# Multiscale Model Within-host and Between-host for Viral Infectious Diseases

**DOI:** 10.1101/174961

**Authors:** Alexis Erich S. Almocera, Van Kinh Nguyen, Esteban A. Hernandez-Vargas

## Abstract

Multiscale models possess the potential to uncover new insights into infectious diseases. Here, a rigorous stability analysis of a multiscale model within-host and between-host is presented. The within-host model describes virus replication and the respective immune response while disease transmission is represented by a simple susceptible-infected (SI) model.

The bridge of within-to between-host is by considering transmission as a function of the viral load of the within-host level. Consequently, stability and bifurcation analyses were developed coupling the two basic reproduction numbers 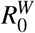 and 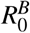for the within- and the between-host subsystems, respectively. Local stability results for each subsystem, such as a unique stable equilibrium point, recapitulate classical approaches to infection and epidemic control.

Using a Lyapunov function, global stability of the between-host system was obtained. A main result was the derivation of the 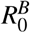 as a general increasing function of 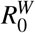. Numerical analyses reveal that a Michaelis-Menten form based on the virus is more likely to recapitulate the behavior between the scales than a form directly proportional to the virus. Our work contributes basic understandings of the two models and casts light on the potential effects of the coupling function on linking the two scales.

## 1 Introduction

Infectious diseases remain a global concern despite the advance of medicine and living conditions [28]. Combating with infectious diseases necessitates multidisciplinary efforts, among which include holistic understandings on the infection mechanisms [12]. Infectious diseases dynamics, however, are governed by many interconnected scales, from complex within-host infection processes, to between hosts, vectors, and environments. In this context, mathematical modelling is an essential tool to untangle the processes, providing understandings of diseases dynamics and public health policies [17].

Typical infectious diseases models [9,25] restrict their dynamics to one of the two scales: *within-host*, focusing on cellular interactions; and *between-host*, focusing on transmission and infection statuses (Figure 1). This restriction allows more analytical tractability [9] but there are problems where it becomes necessary to consider both scales [17,15]. The last decade has witnessed the surfaces of models that bridge infection within- and between-host processes [22,18,11,21,17]. This approach has served as a framework to understand the within-host evolution [1,24], epidemic control and prevention [18,22,23], and beyond [25,15,6]. For example, natural selection of a disease can be investigated by linking pathogen behavior to population dynamics [13]. In [18], the authors investigated the immune responses dynamics in accompany with a SIR model with vaccination. In another study [21], the vector-borne transmission was analyzed with an extended SI model and a within-host subsystem. Applications of multiscale models have broadened the disease modelling landscape and have the potential to accurately describe disease dynamics [25].

**Fig. 1.**
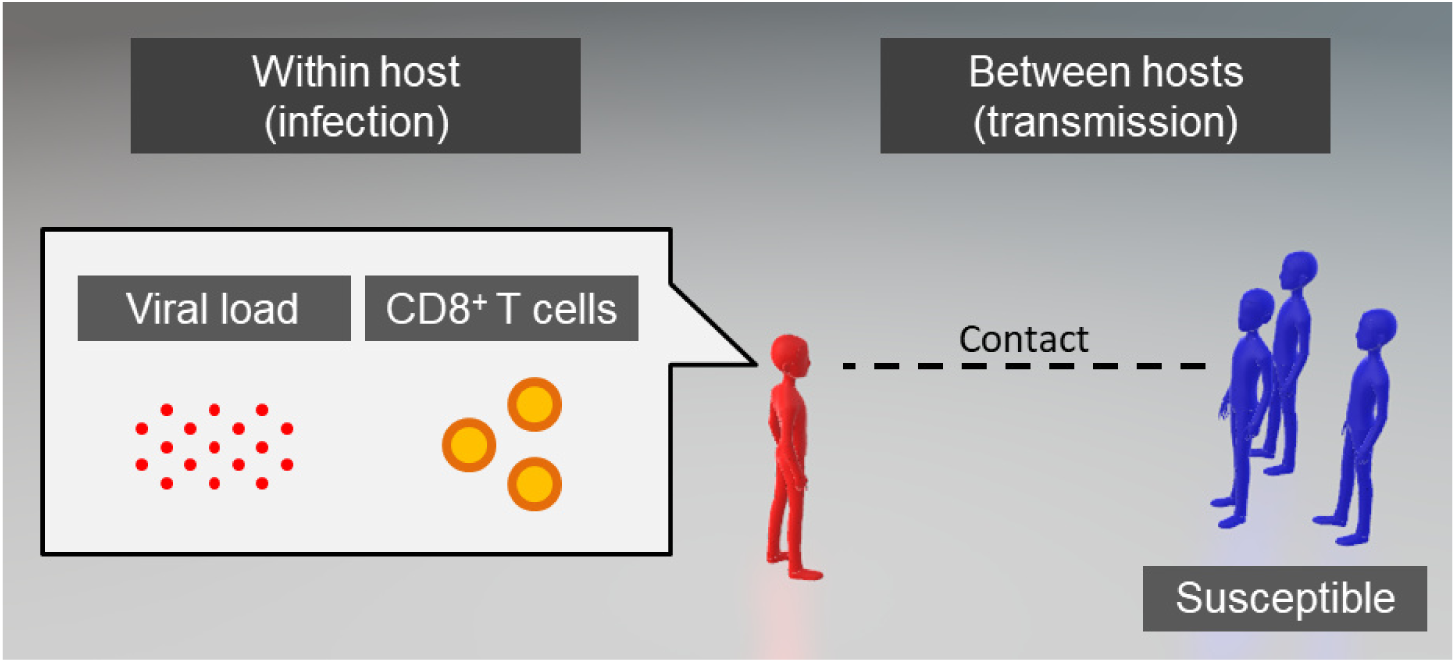
A disease model is usually restricted to one of the following scales: within a host (left) and between hosts (right). Our multiscale model considers both scales, by focusing on the interaction of the virus and immune response within a host, while only considering direct contact between susceptible and infected individuals.

In this paper, we considered a nested model assuming a single coupling direction from within-to between-host. The within-host dynamics is taken from the model formulated by Boianelli et al. [4] whereas between-host transmission dynamics are governed by the susceptible-infective (SI) model [2]. In addition to the quantitative analysis in [4], our results characterize the within-host reproduction number 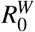 with the existence of a unique positive steady state having viral load *V* ^*\**^ (*i.e.* chronic infections). In addition, we obtain the between-host 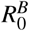assuming the “limiting case” of the between-host subsystem where the viral load remains at the steady state [13]. Results of analytical analyses are accompanied by phase portraits and bifurcation diagrams.

## 2 Materials and Methods

We considered two models that cover the infection dynamics within a host and the transmission between hosts. Each scale is described below with a two-dimensional ordinary differential equation.

### 2.1 Within-host subsystem

We focused on dynamics of virus (*V*) and T-cell populations (*E*) as follows [4]:

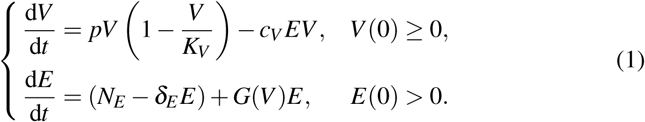

In this model, the virus replicates at a self-limiting rate determined by the intrinsic rate *p* and the carrying capacity *K*_*V*_. The virus is cleared at a per-capita rate proportional to T-cell populations (*c*_*V*_ *E*). In the absence of virus, T cells stay at a homeostatic level given by *E*_0_: = *N*_*E*_/*δ*_*E*_, where *N*_*E*_ and *δ* _*E*_ denote the replenishment rate and half-life of T cells, respectively. Otherwise, T-cells proliferate following a continuous function *G*(*V*) of the viral load. To allow generality for our within-host subsystem, we imposed the following properties:

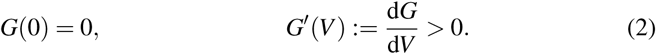

A special case of *G* in the Michaelis-Menten form [4]

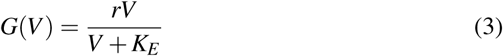

was considered in our analyses.

### 2.2 Between-host subsystem

We considered the Susceptible-Infected (SI) model [2], where the host population is divided into susceptible and infected classes with densities *S* and *I*, respectively. This model reads as

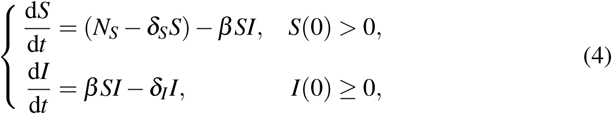

where *β* is the transmission rate. Susceptible individuals entered the population at a rate *N*_*S*_, and both subpopulations have distinct half-life parameters *δ*_*S*_ and *δ*_*I*_. The host population attains the equilibrium size *S*_0_: = *N*_*S*_/*δ*_*S*_ in infection-free situations. For simplicity, we assume all parameters in (4) are fixed.

### 2.3 Bridging scales

Considering experimental observations of the impact of viral load on disease transmission [3,8], we assumed a multiscale model coupling (1) and (4) where *β* = *β* (*V*) has the following properties:

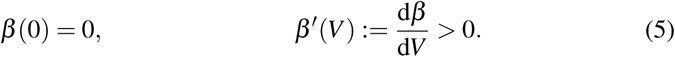

As the exact functional relationship between viral load and the transmission rate can be debatable [15], we considered three different functional forms of the coupling function, including a linear, logistic, and saturation function that satisfies (5) in our numerical analyses. In particular, 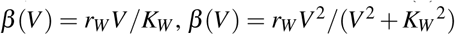, and 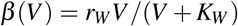[4]. The parameter *r*_*W*_ is the rate of transmission due to the viral load. The parameter *K*_*W*_ is a threshold of the viral load that a host may need to cross to transmit the infection.

### 2.4 Numerical example

To complement our analytical analyses and illustrate infection-transmission dynamics, we obtained numerical results from the two models using the following values:

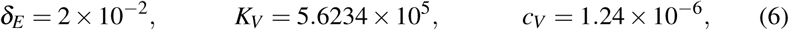

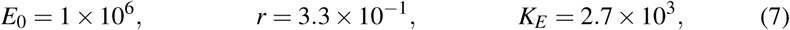

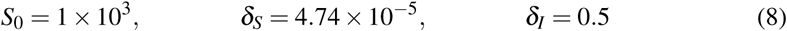

The parameter values in (6) and (7) were taken from [4,19,30], while those in (8) were chosen judiciously. The value of *δ*_*I*_ is based on [5,29], where roughly half of the infected population is removed each day. We assumed that the average life expectancy of a healthy individual is 80 years, hence our choice of *δ*_*S*_.

## 3 Mathematical analysis of the multiscale model

The basic reproduction number (*R*_0_) is used to establish the corner stones for planning public health initiatives, such as the epidemic threshold and herd immunity [2]. The number is defined as the average number of secondary infections caused by a typical infected case in a completely susceptible population over its infectious period [7]. Here, for each of the subsystems, we derived a respective *R*_0_ as a bifurcation parameter of the within-host 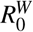 and the between-host 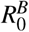, respectively.

### 3.1 Within-host subsystem

The basic reproduction number for the within-host subsystem (1) is given by

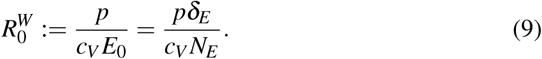

To justify, we have

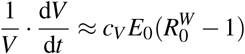

for *V ≈* 0 and *E ≈ E*_0_ (*i.e.* at the outset of infection). Thus, the virus is cleared when 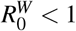 and persists when 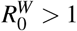. According to the differential equation for *E*, each equilibrium point of (1) takes the form (*V,E*) where *E >* 0. Moreover,

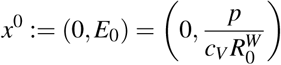

is the unique infection-free equilibrium for (1).

Let *x*^*\**^ be an equilibrium point of (1) whose coordinates are positive. To determine *x* ^*\**^, we define a function *G*_0_ by

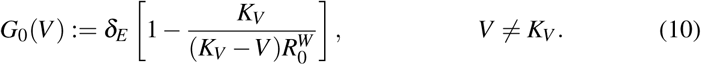

Then the inverse of *G*_0_ is given by

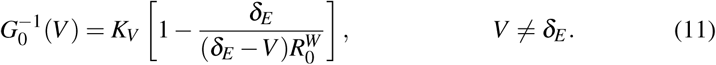

Furthermore, 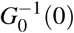 is a zero of *G*_0_, *i.e. 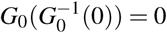*. The following lemma is a major step to obtaining *x*^*\**^.

**Lemma 1** *There exists a solution V of the following equation*

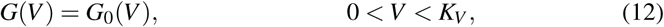

*if and only if 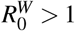. This solution is unique and lies in the open interval* 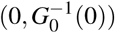.

*Proof* According to the conditions in (2), the function *G* increases, and *G*(*V*) *> G*(0) = 0 for *V >* 0. On the other hand,

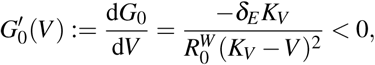

so that *G*_0_ is decreasing. We argue by defining *H* = *G*_0_ *-G*, which is a decreasing function. Then each solution of (12) corresponds to a root of *H* on (0*,K*_*V*_). Note that

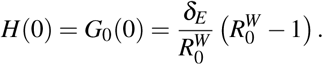

Thus, *H <* 0 on (0*,K*_*V*_) whenever 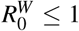. Consequently, *H* has a root on (0*,K*_*V*_), and (12) admits a solution, only if 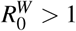.

Now, suppose that 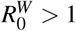1, from which *H*(0) *>* 0 and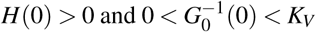. Observe that

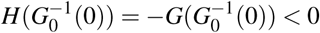

since 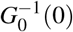 is a root of *G*_0_ and *G >* 0 on (0*,K*_*V*_). Since *H* decreases, *H* has a unique root *v*^*\**^ on 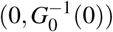 and *H <* 0 on 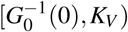. Hence, there exists a solution of (12) if and only if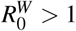; this solution is uniquely given by 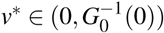. 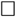

Theorem 1 *Consider the chronic equilibrium point x*^*\**^= (*V*^*\**^, *E*^*\**^)*. Then*

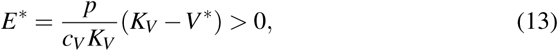

*and V*^*\**^ *is the unique solution of* (12) *in*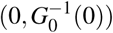 *. Moreover, x*^*\**^ *exists if and only if* 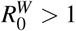.

*Proof* According to the equations of (1), the coordinates of *x*^*\**^satisfy:

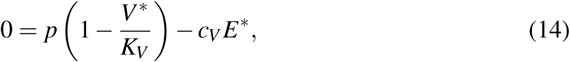

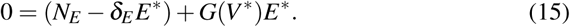

Equation (13) is equivalent to (14), from which 0 *< V*^*\**^ < *K*_*V*_. On the other hand, we transform equation (15) into

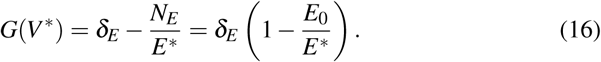

Applying equations (9) and (13) to equation (16), we get

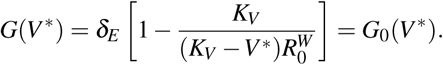

We conclude from Lemma 1 that *x*^*\**^exists, where *V*^*\**^is the unique solution of equation (12) in 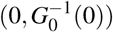, if and only if 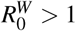. 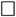

By assuming a specific form of *G*, we can express *V*^*\**^as a function of 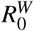. To illustrate, we consider the parameter *r* in equation (3), assume that *r > δ*_*E*_, and introduce the following functions:

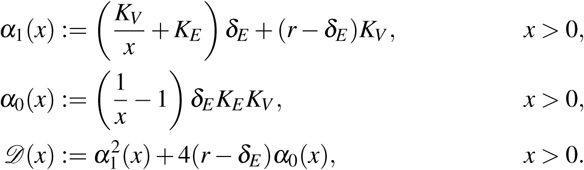

We also let 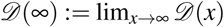. Then we obtain the following:

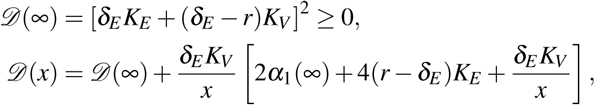

for *x >* 0, where 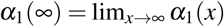. In particular, *D*(*x*) *>* 0; hence, the quadratic polynomial

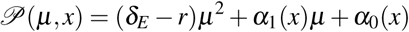

has two distinct roots, the smaller of which is given by

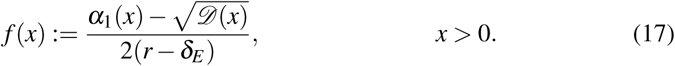

We establish some key properties of *f*.

**Theorem 2** *Assume that r > δ*_*E*_. *Then f is an increasing function (i.e. f* ^*'*^ > 0*). Moreover, f* (*x*) *>* 0 *for x >* 1.

*Proof* We compute the derivative *f* ^*'*^ as

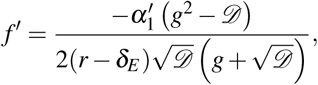

where

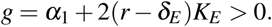

Note that 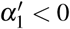 and *D >* 0. Moreover, we obtain

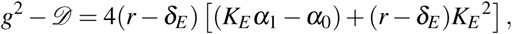

where (*K*_*E*_*α*_1_ *-α*_0_) is a positive constant. Thus, *f* ^*'*^ > 0 and *f* is an increasing function. Finally, we evaluate 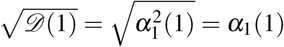, and *f* (1) = 0. Therefore, we have *f* (*x*) *>* 0 for *x >* 1.

The following theorem relates *V*^*\**^ with 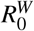 via the function *f*.

Theorem 3 *Assume the inequality r > δ*_*E*_ *and equation* (3)*. If 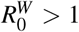, then 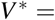 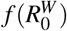*

*Proof* By definition,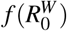 is the smaller root of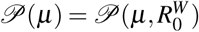 in *µ*. Denoting the larger root by *p*_(+)_, i.e. 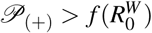. Appealing to Theorem 1, and applying (3) to (12), we obtain *V*^*\**^as a root of *P*(*µ*) in 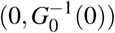. Thus, either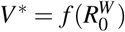 or *V*^*\**^= *P*_(+)_.

Recalling that 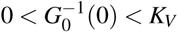whenever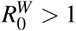, we compute

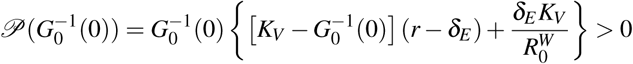

and note that *P*(*µ*) is concave down. Thus,

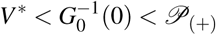

and we must have 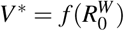.

We establish local stability of (1) as follows.

**Theorem 4** *The following statements hold:*

1. *The infection-free equilibrium point x*^0^ = (0*,E*_0_) *is*
  a. *asymptotically stable if 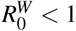*
  b. *nonhyperbolic if 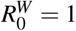, and*
  c. *a saddle point if 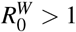.*
2. *If 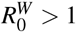, then the equilibrium point x*^*\**^ = (*V*^*\**^, *E*^*\**^) *is asymptotically stable.*

*Proof* Consider the following Jacobian matrix associated with (1):

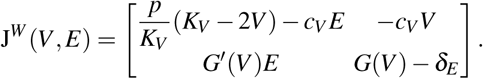

Then J^*W*^ (*x*^0^) is a lower-triangular matrix. Moreover, recalling (2) and (9), we obtain the following eigenvalues of J^*W*^ (*x*^0^):

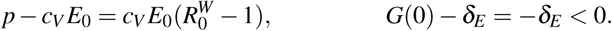

This establishes statement 1.

Now, we assume 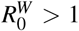 and appeal to Theorem 1. Denote the (*k,k*)-entry of J^*W*^ (*x*^*\**^) by 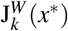. Then equation (13) yields

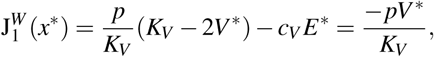

while equations (10) and (12) (where *V* = *V*^*\**^) yield

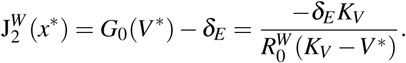

Thus, J^*W*^ (*x*^*\**^) has a negative trace. On the other hand, *G*^*'*^(*V*^*\**^) *>* 0 by (2), from which J^*W*^ (*x*^*\**^) has a positive determinant. Accordingly, each eigenvalue of J^*W*^ (*x*^*\**^) has a negative real part. Therefore, *x*^*\**^is asymptotically stable. Statement 2 is proven.

Theorem 4 indicates a unique equilibrium point 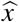for the system (1) that is either asymptotically stable or is nonhyperbolic; that is,

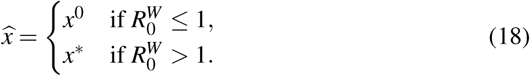

We need the following hypothesis in our forthcoming analysis of the between-host subsystem (4):

(H) For each solution of (1) with *V* (0) *>* 0 and *E*(0) *>* 0, we have

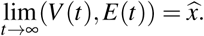

Thus, we say that 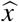 is *globally asymptotically stable*.

A consequence for those solutions in hypothesis (H) is that

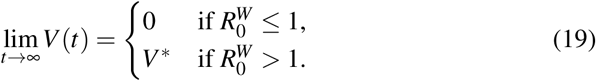

This reiterates 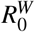 as an infection threshold for the system (1): the virus does not survive when 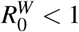, and persists when 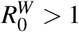. The phase plane portraits in Figures 2 supports the global stability of 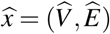 in hypothesis (**H**).

**Fig. 2.**
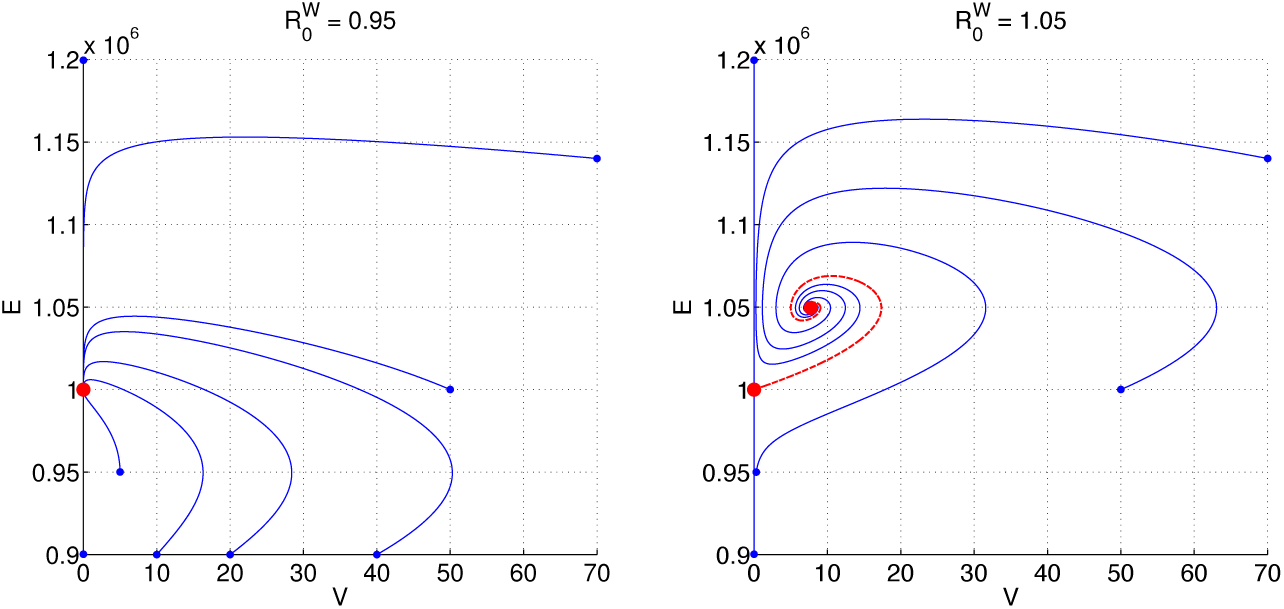
Phase plane portraits for the within-host subsystem (1) at two different values of 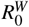 Each curve represents a solution (*V* (*t*)*,E*(*t*)) for *t ≥* 0, with initial time indicated by the blue endpoint. The red curve represents the heteroclinic orbit that connects *x*^0^ to *x*^*\**^. Red points indicate equilibrium points.The following parameter values were used: *c*_*V*_ = 1.24 *×* 10^*-*6^; *N*_*E*_ = 2 *×* 10^4^; *δ*_*E*_ = 2 *×* 10^*-*^2; *r* = 3.3 *×* 10^*-*^1;*K*_*E*_ = 2.7 *×* 10^3^; *K*_*V*_ = 5.6234 *×* 10^5^; and 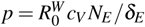

### 3.2 Coupled within-host and between-host model

Consider hypothesis (H), and assume that 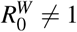. Then, the system (1) equilibrates to 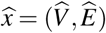, where

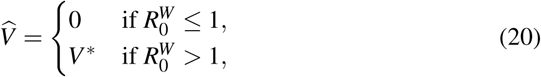

by (18). Thus, we may assume in (4) that 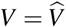, yielding the following system:

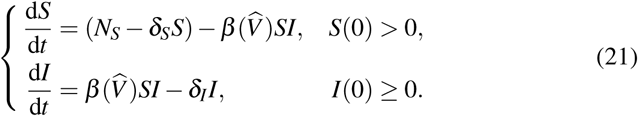

Let us call (21) the *limiting case* of the between-host subsystem.

We focus on the case where, 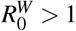. To justify, suppose that 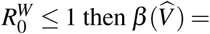 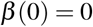 by (20), and the equations in (21) yield lim_*t*→∞_(*S*(*t*)*,I*(*t*)) = (*S*_0_,0) regardless of initial condition and parameter values. If 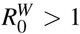, then (21) is rewritten as follows:

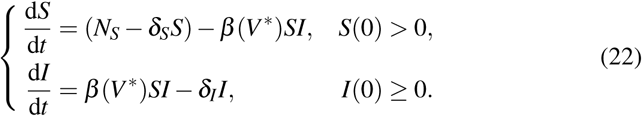

We assume that 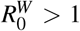 and analyze the limiting case (22). We realize our basic reproduction number for the limiting case (22) of the between-host subsystem as

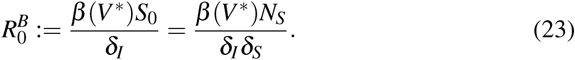

Indeed, from (22) we have 1

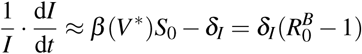

for *S ≈ S*_0_ and *I ≈* 0, i.e. in a disease-free population. Thus, we expect the disease to be eradicated when 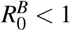, and an endemic to occur when 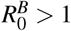.

It follows from the differential equation for *S* that each equilibrium point of the system (22) is of the form (*S,I*), where *S >* 0. Moreover,

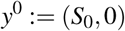

is the unique disease-free equilibrium point for (22).

Let *y*^*\**^= (*S*^*\**^,*I*^*\**^) be an endemic equilibrium point of (22), where both *S*^*\**^and *I*^*\**^are positive. Then *y*^*\**^is unique, according to the following theorem.

**Theorem 5** *Assume that 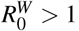. Then y*^*\**^= (*S*^*\**^,*I*^*\**^) *exists, with unique coordinates*

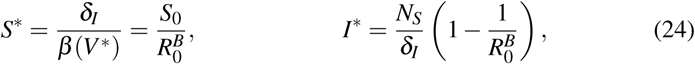

*if and only if 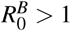.*

*Proof* From the equations in (22), we obtain the following:

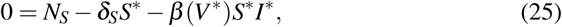

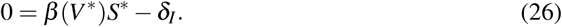

Equation (26) gives the desired expression for S* Thus from equation (25), we have the following:

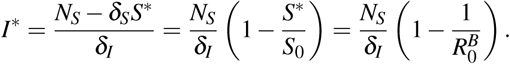

Thus, we obtain (24). Consequently, *y*^*\**^exists if *I*^*\**^ *>* 0, i.e. 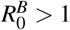.

It is possible for 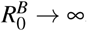 representing an increased transmission of the disease. In this case, it follows from equation (24) that *S*^*^ ⟶ ∞ all the population at the endemic steady state is infected.

The local stability for the system (22) is given by the following result.

Theorem 6 *Assuming that 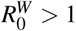, the following statements hold:*

1. *The disease-free equilibrium point y*^0^ *is*
  a. *asymptotically stable if 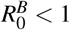,*
  b. *nonhyperbolic if 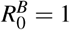, and*
  c. *a saddle point if 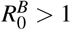.*
2. *If 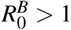, then the endemic equilibrium point y*^*\**^ = (*S*^*\**^,*I*^*\**^) *is asymptotically stable.*

*Proof* We begin with the following Jacobian matrix associated with (22):

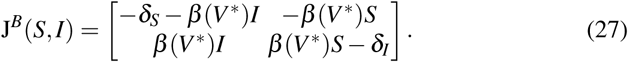

Then J^*B*^(*y*^0^) is an upper-triangular matrix, with eigenvalues *-δ*_*S*_ and

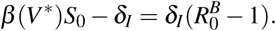

Thus, statement 1 holds.

To prove statement 2, we assume that 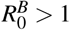 and denote the (*k,k*)-entry of J^*B*^(*y*^*\**^) by 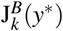 Note that *β* (*V*^*\**^) *>* 0 by (5), hence 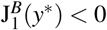. Meanwhile,

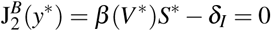

by Theorem 5. Thus, the matrix J^*B*^(*y*^*\**^) has a negative trace and a positive determinant, and each eigenvalue of J^*B*^(*y*^*\**^) has a negative real part. Therefore, statement 2 holds.

We obtain from Theorem 6 a unique equilibrium point 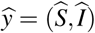 of (22) that is either asymptotically stable or nonhyperbolic. That is,

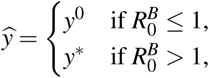

from which 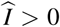 if and only if 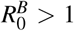 We establish the global stability of 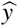 follows.

**Theorem 7** *Assuming that 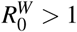, then*

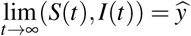

*for each solution of* (22)*, such that I*(0) *>* 0 *whenever 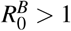.*

*Proof* Let us define the following sets

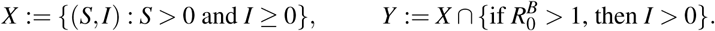

Denote by 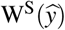 the intersection of *X* with the stable manifold of 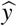. It is enough to establish

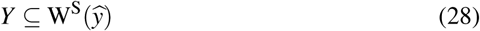

to prove the theorem. We remark that 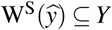 due to the forward invariance of the *I* = 0 plane.

We appeal to the LaSalle invariance principle [20,31] to establish (28). For *w* = *S,I*, we define a function *ℒ*_*w*_: *Y* ⟶ *R* by

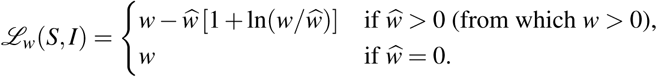

Consider the sum *ℒ* = *ℒ*_*S*_ + *ℒ*_*I*_ and its time derivative *ℒ*^*'*^. Let

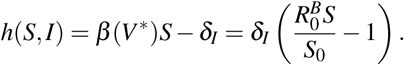

Then we have

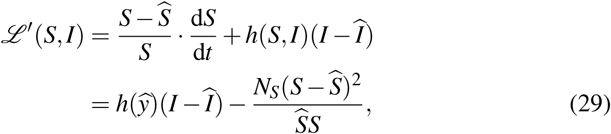

after some computations. Observe that

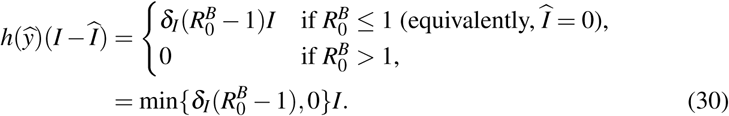

Substituting (30) to (29) yields

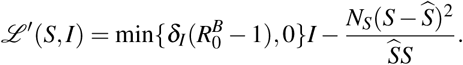

Thus, *ℒ* is a Lyapunov function, and *ℒ*^*'*^(*S,I*) = 0 if and only if the following property holds: 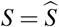, and 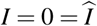 whenever 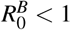.

Now, consider the following set:

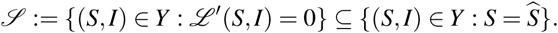

Then each solution taking values in *S* satisfies

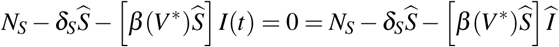

and 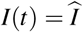 for all *t* ≥ 0; that solution is necessarily 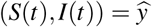. Consequently, 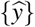 is the largest invariant subset of *S*. Therefore, equation (28) holds by the LaSalle invariance principle.

In addition to Theorem 7, the forward invariance of the *I* = 0 plane yields lim_t→∞_(S(t); I(t))= (S0;0)=y^0^ (regardless of 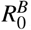 whenever *I*(0) = 0. Figures 3 illustrates the phase plane portraits of the system (22), showing the global stability of 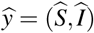 established in Theorem 7.

**Fig. 3.**
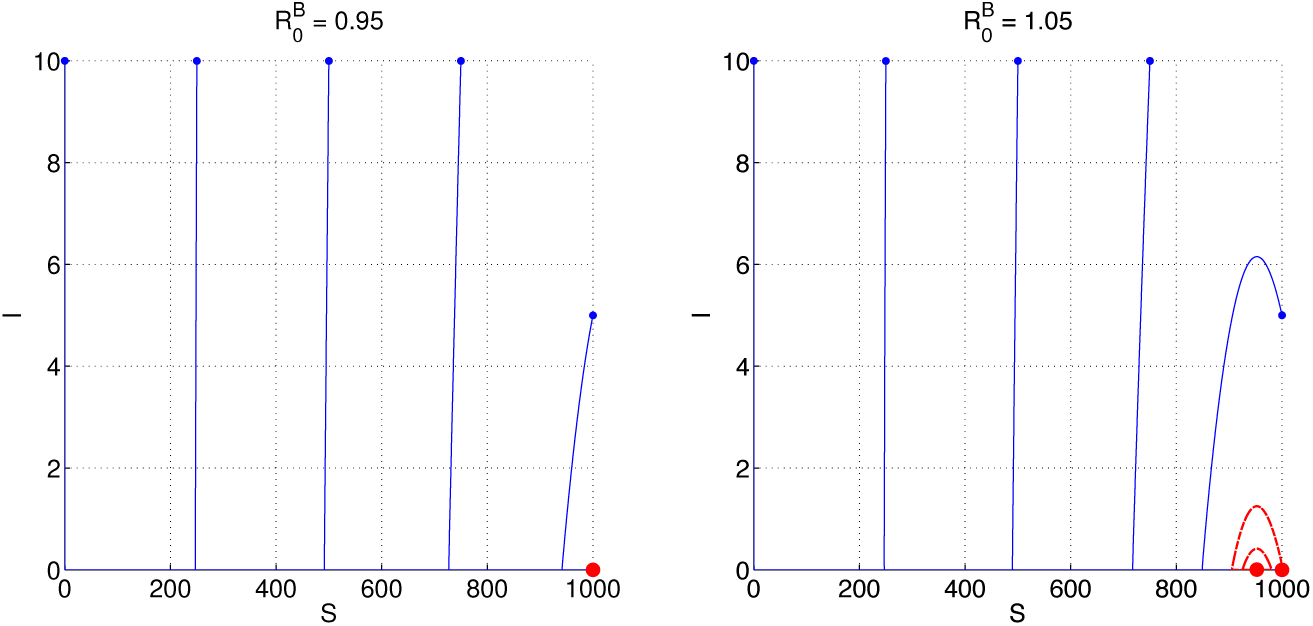
Phase plane portraits for the between-host subsystem (22) at two different values of 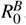. Each curve represents a solution (*S*(*t*)*,I*(*t*)) for *t ≥* 0, with initial time indicated by the blue endpoint. The red curve represents the heteroclinic orbit that connects *y*^*\**^to *y*_0_. Red points indicate equilibrium points. The simulation reveals a slow manifold near the *S*-axis, to which each blue-colored solution curve is asymptotic. The following values were used: 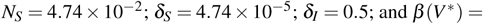 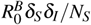

### 3.3 Relating the two basic reproduction numbers

It is useful to describe the dynamics of both systems (1) and (22) with respect to a single basic reproduction number. Assuming that 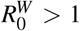, we have

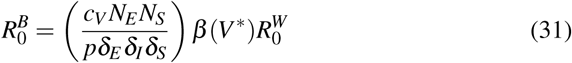

by equations (9) and (23). Here, we do not consider the dependence of *V*^*\**^on 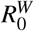 With further assumptions, we establish below that 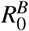 is an increasing function of 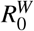.

**Theorem 8** *Assuming that r > δ*_*E*_, *define a function F on* (0, ∞) *by*

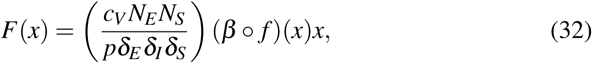

*where f is given by equation* (17)*. Then F*^*'*^(*x*) *>* 0 *for x >* 1*. Furthermore, if equation* (3) *holds, then*

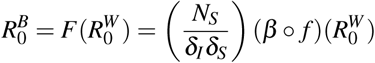

*For 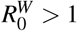.*

*Proof* For *x >* 1, we have

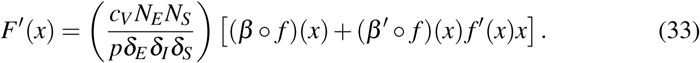

Observe from (5) and Theorem 2 that *β*, *β*^*'*^, and their derivatives are positive on the interval (1,∞). Thus, we have *F*^*'*^(*x*) *>* 0. Under the assumption of (3), Theorem 3 yields 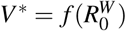 for 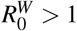, from which 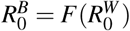 by equation (31).

Figure 4 depicts the relationship between basic reproduction numbers, given by the equation 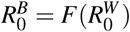 in Theorem 8. Here, three examples for *β* are considered. Aside from validating the increasing property of *F*, this figure shows that the relationship depends on the definition of *β*.

## 4 Discussion

Multiscale models have shown capable to generate hypotheses and new insights into infectious disease mechanisms [15]. Linking the two scales has led to the identification of missing pieces in infectious disease research and is the only way to capture effects of the feedback between two scales [24]. While there exist many obstacles in the approach [15,24,14], multiscale modeling is a promising approach that would improve our understandings on the epidemic control and prevention.

**Fig. 4.**
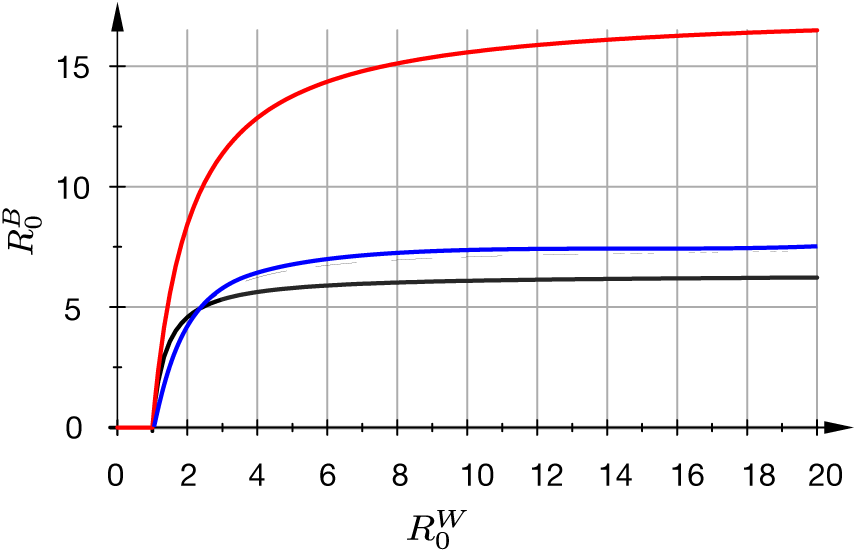
The relationship between the two basic reproduction numbers, as given by the equation 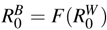 in Theorem 8. This relationship depends on the definition of the transmission rate: *β* (*V*) = *r*_*W*_*V /*(*V* + *K*_*W*_) (black curve); *β* (*V*) = *r*_*W*_*V* ^2^*/*(*V* ^2^ + *K*_*W*_ ^2^) (blue curve); and *β* (*V*) = *r*_*W*_*V /K*_*W*_ (red curve). In all examples, *r*_*W*_ = 0.005 and *K*_*W*_ = 100.

In this paper, we analyzed a multiscale model that comprises a within-host viral infection model, nested in a simple SI transmission model. A rigorous stability analysis and numerical simulations for the multiscale model were performed. We derived stability analyses considering that the stimulation of the immune system by the virus (*V*) is represented by any function *G*(*V*) with the properties *G*(0) = 0 and *G*^*'*^(*V*) *>* 0.

With a specific expression for *G*(*V*), we obtained an expression 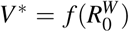 which consequently provides a relationship between the basic reproduction numbers. We conjecture that the existence of *f* holds in general. Meanwhile, the global stability of the between-host system (Theorem 7) was established through the construction of a Lyapunov function. We derived two basic reproduction numbers,

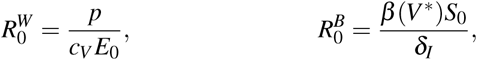

for the within-host level and between-host, respectively. Viral infections with 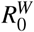 below unity would derive to clear the infection within a host; otherwise, a chronic state could be established (Figure 2). For viruses with a very high replication rate, such as avian influenza [16] and Ebola virus [26], the virus replication likely outpaces the developing immune response [16,27]. Early interventions to stem down viral replication rate could promote survival and recovery from infection [16,27]. Notable examples are the use of neuraminidase inhibitors in containing influenza infection [22,10] and the use of antiretroviral treatment (ART) as a public prevention measure for HIV [17].

Furthermore, given the same disease and similar intervention approaches are in-place, i.e. *δ*_*I*_ and *β* (*V*^*\**^) are fixed, the epidemic size then depends on the influx of new susceptible individuals (Equation (24)). Approaches, such as childhood vaccinations, diverge the influxes into non-susceptible classes and thus could lead to a disease eradication. As per the SI model's assumption of a constant population, equation (24) also shows that the epidemic size will saturate regardless of 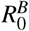. As 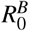 will also saturate as 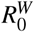 crosses a certain value (Figure 4), it follows that epidemic interventions aim at controlling viral replication would need to be highly effective to influence the epidemic size. In other words, if an intervention can largely reduce the 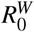 but this value is still above a certain threshold then the intervention would have a negligible impact on the epidemic size. For our set of parameter values (Section 2.4), this threshold is approximately 1.5 (Figure 5).

**Fig. 5.**
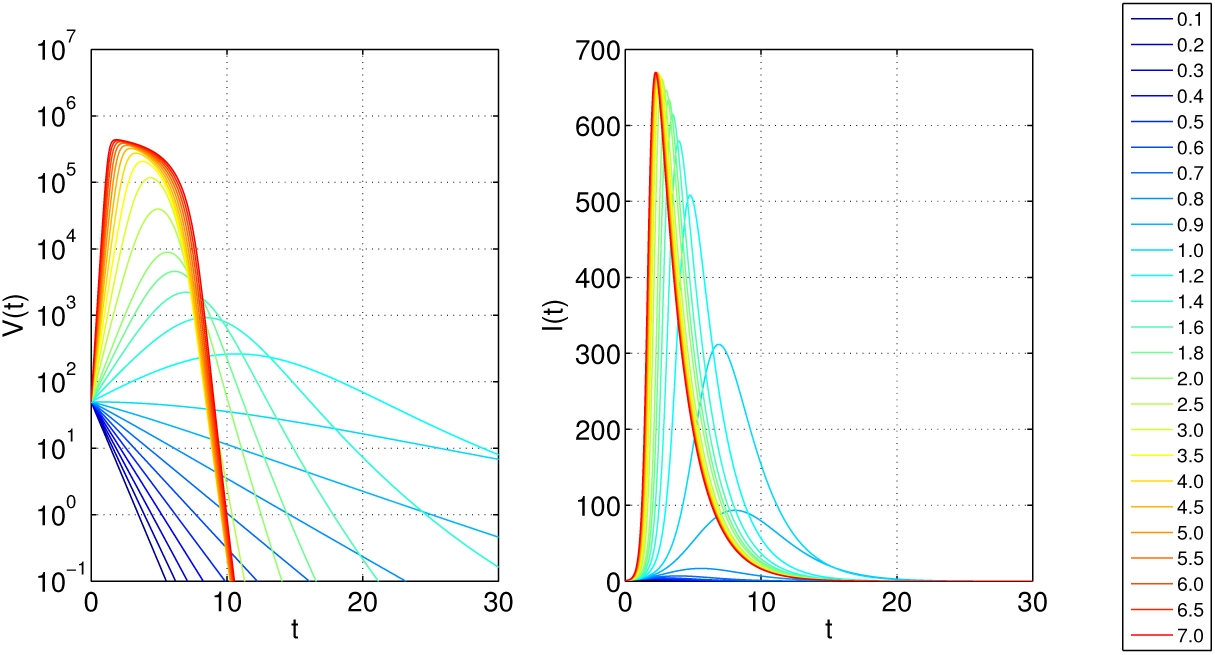
The trajectories *V* (*t*) and *I*(*t*) depend on the within-host reproduction number. Each curve denotes the same solution, colored according to a unique value of 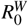; see the legend on the right. The saturation function *β* (*V*) = *r*_*W*_*V /*(*V* + *K*_*W*_) was used, where *r*_*W*_ = 0.005 and *K*_*W*_ = 100. The initial conditions are *V* (0) = 50, *E*(0) = 10^6^, *S*(0) = 999, and *I*(0) = 1.

**Fig. 6.**
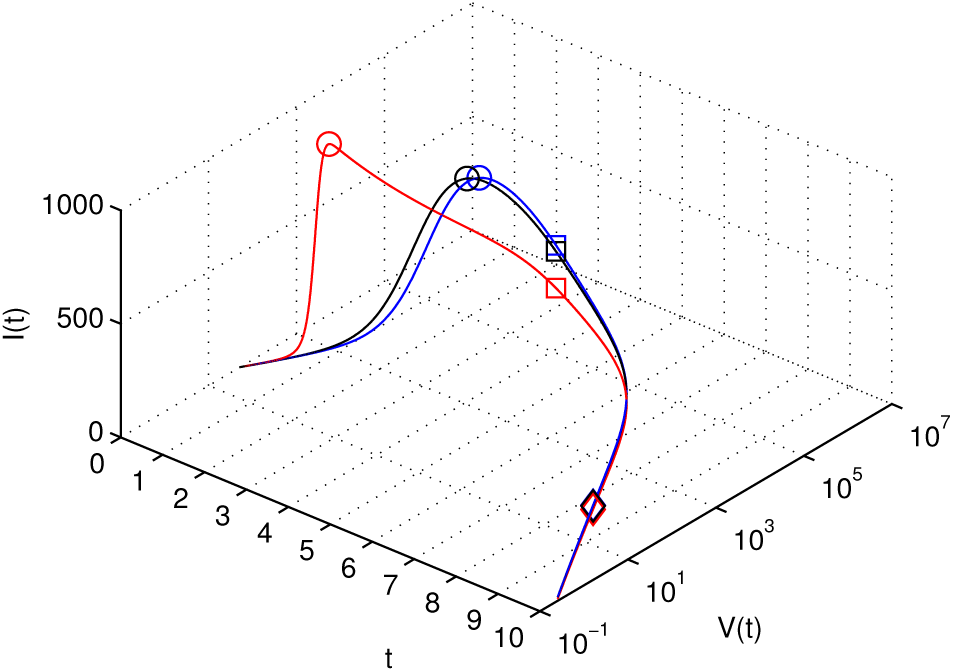
The solution trajectory of the multiscale model depends on the definition of the transmission rate *β* (*V*). The same examples of *β* (*V*) are considered as in Figure 4: saturation (blue), linear (red), and logistic (black). In each case, the solution describes a peak in the infected cases (circle), followed by a peak in the viral load (square), before the viral load reduces to less than the detectable threshold of 50 PFU/mL (diamond). The initial conditions are *V* (0) = 50, *E*(0) = 10^6^, *S*(0) = 999, and *I*(0) = 1.

The above results could still hold when the coupling function between the two scales was varied (Figure 4). The correct link function for many diseases is still debatable because empirical data are lacking [24,15,14]. For this reason, we tested three functional forms expressing the transmission rate based on viral load including a linear, logistic, and Michaelis-Menten function as discussed elsewhere [15]. Figure 4 shows that while the effect of 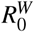 on 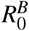 is qualitatively indifferent among the functions, there is a marked difference quantitatively. Linearly coupling led to the transmission increased roughly triple compared to the others. Thus, coupling using a linear function for a pathogen may lead to an overestimation of a disease transmission. This is illustrated in more details in Figure 6, for the same set of parameter, the linear function would lead to a much faster and stronger epidemic trajectory.

The analysis of the dynamics from within-to between-host of infectious disease requires the coupling of systems that are usually studied in isolation. The high level of detail involved in this approach entails greater efforts. As of now a complete framework to study multiscale infectious diseases has not prevailed, but it is a constant target. This work contributes basic understandings of the two models in considered and casts light on potential effects of the coupling function on linking the two scales. Future experimental studies are needed to verify our results to improve multiscale models within-host and between-host.

